# CrisprBuildr: an open-source application for CRISPR-mediated genome engineering in *Drosophila melanogaster*

**DOI:** 10.1101/2025.02.28.640916

**Authors:** Nicole Horsley, Adam von Barnau Sythoff, Mark Delgado, Selina Liu, Clemens Cabernard

## Abstract

CRISPR/Cas9 is a powerful tool for targeted genome engineering experiments. With CRISPR/Cas9, genes can be deleted or modified by inserting small peptides, fluorescent proteins or other tags for protein labelling experiments. Such experiments are important for detailed protein characterization *in vivo*. However, designing and cloning the corresponding constructs can be repetitive, time consuming and laborious. To aid users in CRISPR/Cas9-based genome engineering experiments, we built CrisprBuildr, a web-based application that allows users to delete genes or insert fluorescent proteins at the N- or C-terminus of their gene of choice. The application is built on the *Drosophila melanogaster* genome but can be used as a template for other available genomes. We have also generated new tagging vectors, using EGFP and mCherry combined with the small peptide SspB-Q73R for use in iLID-based optogenetic experiments. CrisprBuildr guides users through the process of designing guide RNAs and repair template vectors. CrisprBuildr is an open-source application and future releases could incorporate additional tagging or deletion vectors, genomes or CRISPR applications.

## Introduction

Genome engineering, such as deleting genes, introducing point mutations or inserting defined DNA sequences at specific loci, has been revolutionized with the advent of CRISPR/Cas9 (Sternberg et al. 2014). The ability to precisely manipulate the genome has significantly enhanced the molecular genetic toolkit from bacteria to human cells. CRISPR/Cas9 positively impacts basic research with classic model organisms such *Drosophila melanogaster*. ∼ 62% of human disease genes are conserved in *Drosophila* (Adams et al. 2000; Rubin et al. 2000), and loss-of-function and protein localization studies aid in the functional characterization of genes and molecular processes.

Visualizing protein localization with high spatial and temporal dynamics is a key tool to understand protein function. Fusing proteins of interest with fluorescent proteins (FP) for live cell analysis has become a powerful tool to study protein function in various cellular contexts. Of particular interest are genetically encoded FP fusions since they do not need to be transfected for each experiment (Kanca et al. 2017). Genetically encoded fusion proteins are usually inserted in random locations in the genome. Their expression is either controlled via endogenous promotors or enhancers, usually cloned upstream of the fusion protein, or through binary expression systems. For instance, the Gal4/UAS system has been used extensively to express FP-fusions under the control of tissue-specific Gal4 lines in *Drosophila melanogaster* (BRAND and Perrimon 1993) as well as zebrafish (Scheer and Campos-Ortega 1999).

With the discovery of CRISPR/Cas9 based genome engineering methods, it is now possible to integrate FPs with high precision, replacing unlabeled proteins with genetically encoded protein-FP fusions (Bier et al. 2018). CRISPR/Cas9 creates double-stranded breaks (DSBs) at defined positions in the fly genome by guiding the endonuclease Cas9 with a programmable guide RNA (gRNA) (Gratz, Wildonger, et al. 2013; Gratz, Cummings, et al. 2013; Gratz et al. 2014; Gratz et al. 2023a; Gratz et al. 2023b). Such DSBs can either be repaired using nonhomologous end joining (NHEJ) or homology-directed repair (HDR) pathways. For the purpose of endogenous gene tagging, the HDR pathway is preferred, which requires single or double-stranded exogenous DNA as repair templates (Bier et al. 2018). Designing CRISPR-based genome engineering experiments can be performed routinely in labs with basic experience in molecular biology and step-by-step protocols and web-resources have been published previously (Bier et al. 2018; Gratz et al. 2023a). However, planning and designing constructs for CRISPR experiments can be challenging, time consuming and laborious. Here, we report the generation of a new set of vectors and a web-based cloning tool, called CrisprBuildr, to aid in the planning and cloning of repair plasmids for generating deletion alleles or for tagging proteins of interest at the endogenous locus with FPs. CrisprBuilder is a bricolage of existing webtools, assembled into a step-by-step workflow, guiding users through the deletion or tagging process *in-silico*. CrisprBuildr reduces the time needed to plan and design Crispr-based genome engineering experiments. The tool also provides a platform to train new users on CRISPR-based genome engineering experiments. CrisprBuilder is built on an open platform, and we envision it to be extended by including other tagging vectors, genomes or CRISPR-based applications.

## Results

### Construction of new protein tagging vectors for protein localization and mislocalization experiments

We constructed a new set of cloning vectors with the goal to (1) tag genes of interest at their endogenous locus for localization studies and (2) combine it with the optogenetic iLID (Guntas et al. 2015) system for protein mislocalization experiments. To this end, we modified the previously published pHD-DsRed-attP vector (Gratz et al. 2014) (Figure 1A), inserting a cassette containing SspB(R73Q) and a fluorescent protein (FP; EGFP or mCherry) upstream of DsRed for either N or C-terminal tagging. SspB(R73Q) is a 13kD adaptor protein from *E. Coli* with improved binding affinity to the *E. Coli* protein SsrA (Guntas et al. 2015). pHD-SspB::FP-DsRed, pHD-FP::SspB-DsRed vectors generate fusion proteins compatible with the optogenetic iLID system (Guntas et al. 2015), which utilizes 7 residues of SsrA, inserted into the photoactive LOV2 element of Avena sativa (AsLOV). SspB is cloned in frame with FPs and either located at the very N- or C-terminus to be accessible for iLID’s SsrA residue (Figure 1B, C). We previously demonstrated the utility of the iLID system *in vivo* by trapping centrosomal proteins to centrioles in dividing fly neuroblasts (Gallaud et al. 2020). We retained the two LoxP sites of pHD-DsRed and used the remaining LoxP site as a peptide linker between the protein of interest and the SspB-FP cassette after Cre-mediated site-specific recombination (Figure 1D, E). The two multiple cloning sites (MCSs) are used to clone in the homology arms for the desired locus.

**Figure 1:**
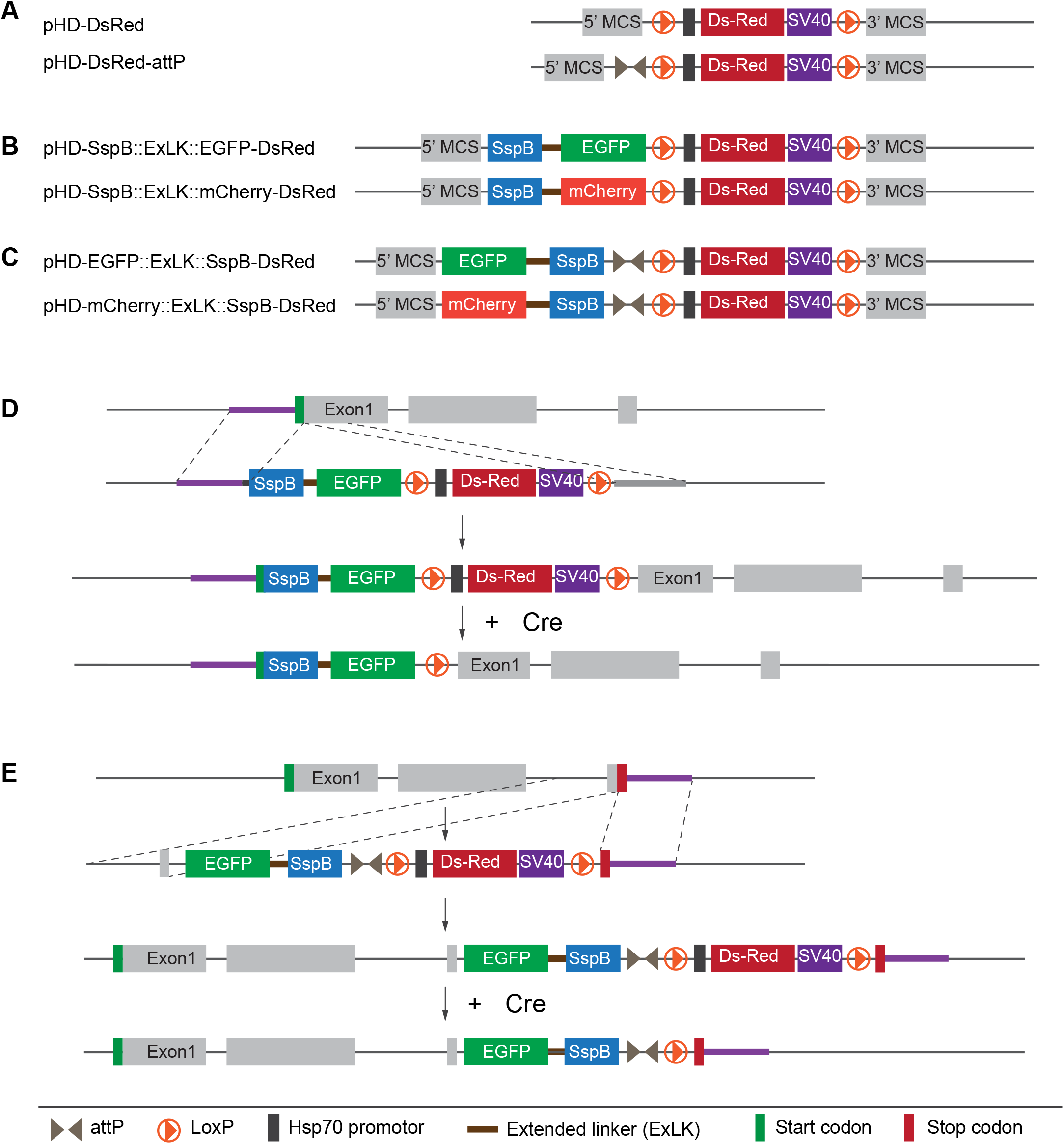
Tagging vectors for Crispr/Cas9 genome engineering. **(A)** Existing vectors used for creating gene deletions. New vectors for **(B)** N-terminal or **(C)** C-terminal tagging experiments. The vectors allow for the insertion of SspB::EGFP or SspB::mCherry cassettes for iLID-mediated optogenetic experiments and protein localization studies. **(D)** After inserting the SspB::EGFP/mCherry at the N-terminus or **(E)** the mCherry/EGFP::SspB at the C-terminus, the DsRed cassette can be floxed out using the existing LoxP sites. The resulting fusion proteins contain the SspB at the end of the protein for better accessibility.

To demonstrate the utility of these vectors, we inserted the SspB::EGFP or EGFP::SssB into the N-terminus or C-terminus of the Par complex component atypical Protein Kinase C (aPKC; Protein Kinase C iota in vertebrates), AuroraB (AurB; AurK in vertebrates) and Myosin’s regulatory light chain Spaghetti Squash (Sqh; MYL9 in vertebrates). To this end, we cloned in homology arms of the corresponding genes into the modified pHD-SspB:::EGFP-DsRed-attP or pHD-EGFP::SspB-DsRed-attP and injected the plasmids together with guide RNAs into Cas9 expressing embryos. After floxing out the DsRed cassette of successful knock-in lines, we imaged third instar larval neuroblasts expressing these fusion proteins together with the microtubule marker cherry::Jupiter (Karpova et al. 2006; Cabernard and Doe 2009).

In mitotic fly neuroblasts, aPKC is localized on the apical neuroblast cortex (Wodarz et al. 2000; Gallaud et al. 2017), and our SspB::EGFP::aPKC line reproduced this localization pattern (Figure 2A). AurB is a member of the chromosome passenger complex (CPC) and both SspB::EGFP::AurB and AurB::EGFP::SspB correctly localized to the centromeric region as well as the ingressing cleavage furrow similar to other CPC components (Roth et al. 2015) (Figure 2B, C). Finally, Sqh::EGFP::SspB is transitioning from a cortical localization, with transient apical enrichment, to the ingressing cleavage furrow as previously described with other Sqh::EGFP transgenic lines (Royou et al. 2002; Barros et al. 2003; Cabernard et al. 2010; Roubinet et al. 2017; Tsankova et al. 2017) (Figure 2D). We conclude that the insertion of the SspB::EGFP or EGFP::SspB at the endogenous locus results in fusion proteins with accurate localization dynamics in fly neural stem cells.

**Figure 2:**
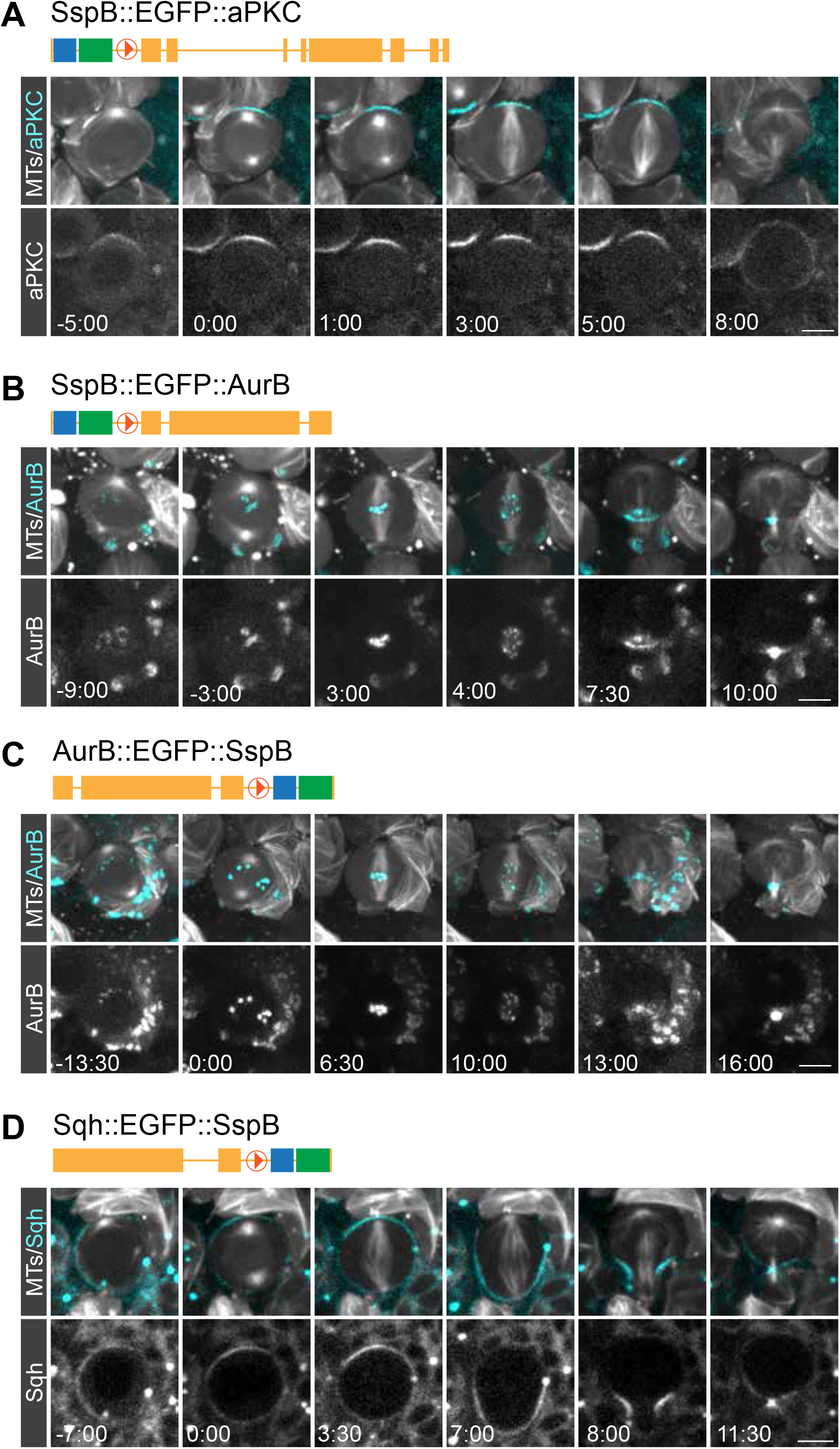
Example fly lines expressing fusion proteins in fly neuroblasts. **(A)** N-terminal tagging of aPKC with SspB::EGFP. aPKC correctly localizes on the apical neuroblast cortex throughout mitosis. N-termianl **(B)** and **(C)** C-termial tagging of AuroraB with SspB::EGFP and EGFP::SspB, respectively. Both lines show AurB localized on kinetochores and the cleavage furrow in telophase. **(D)** C-terminal tagging of the Myosin’s regulatory subunit Spaghetti Squash (Sqh) with EGFP::SspB. Sqh::EGFP correctly localizes at the neuroblast cortex throughout mitosis. Scale bar is 5μm; Time in mins:secs

### CrisprBuildr, an open-source customizable web application for designing gene deletion and tagging constructs

Next, we sought to streamline the generation of CRISPR-based genome engineering experiments. To this end, we built a custom-made cloning app, called CrisprBuildr http://142.93.118.6/. CrisprBuildr aids users in the design of (1) deletion and (2) tagging experiments for genes of their choice. Specifically, CrisprBuildr allows users to generate cloning maps for deletion or tagging experiments. CrisprBuildr accepts Flybase gene symbols or Flybase gene IDs. CrisprBuildr connects to Flybase (www.flybase.org) to retrieve the locus information of the gene of interest. Users can choose to design deletion or tagging experiments after inserting the gene of interest in the search field. In both cases, the desired isoform can be selected from a pull-down list. For tagging experiments, users are prompted to select N-vs. C-terminal tagging and based on this decision, CrisprBuildr will search around the start or stop codon for suitable target sites to induce double-stranded breaks. To this end, CrisprBuildr is interacting with targetfinder.flycrispr.neuro.brown.edu/ to find suitable Crispr sites and subsequently cross checks the sites’ efficiency by connecting to www.flyrnai.org/evaluateCrispr/. After selecting the best Crispr cut sites, CrisprBuilder will search for suitable primer sites within 400-600 bases and 1000-1200 bases both upstream and downstream from the target area to search for suitable amplification and sequencing primers. For CRISPR tagging experiments we aim to include ∼ 1000bp homology arms (see Figure 1D, E) flanking the desired insertion site but shorter arms can also be used (Kanca et al. 2019). CrisprBuildr provides the corresponding amplification and sequencing primers. It will submit this information via simulated browser to bioinfo.ut.ee/primer3-0.4.0/ and will return the selection to the user. As a last step, users can choose a pHD-SspB::FP-DsRed (for N-terminal tagging) or pHD-FP::SspB-DsRed (for C-terminal tagging), containing a fluorescent protein of their choice. We designed vectors containing either an EGFP (Cormack et al. 1996) or mCherry (Shaner et al. 2004) (Figure 3A, B).

**Figure 3:**
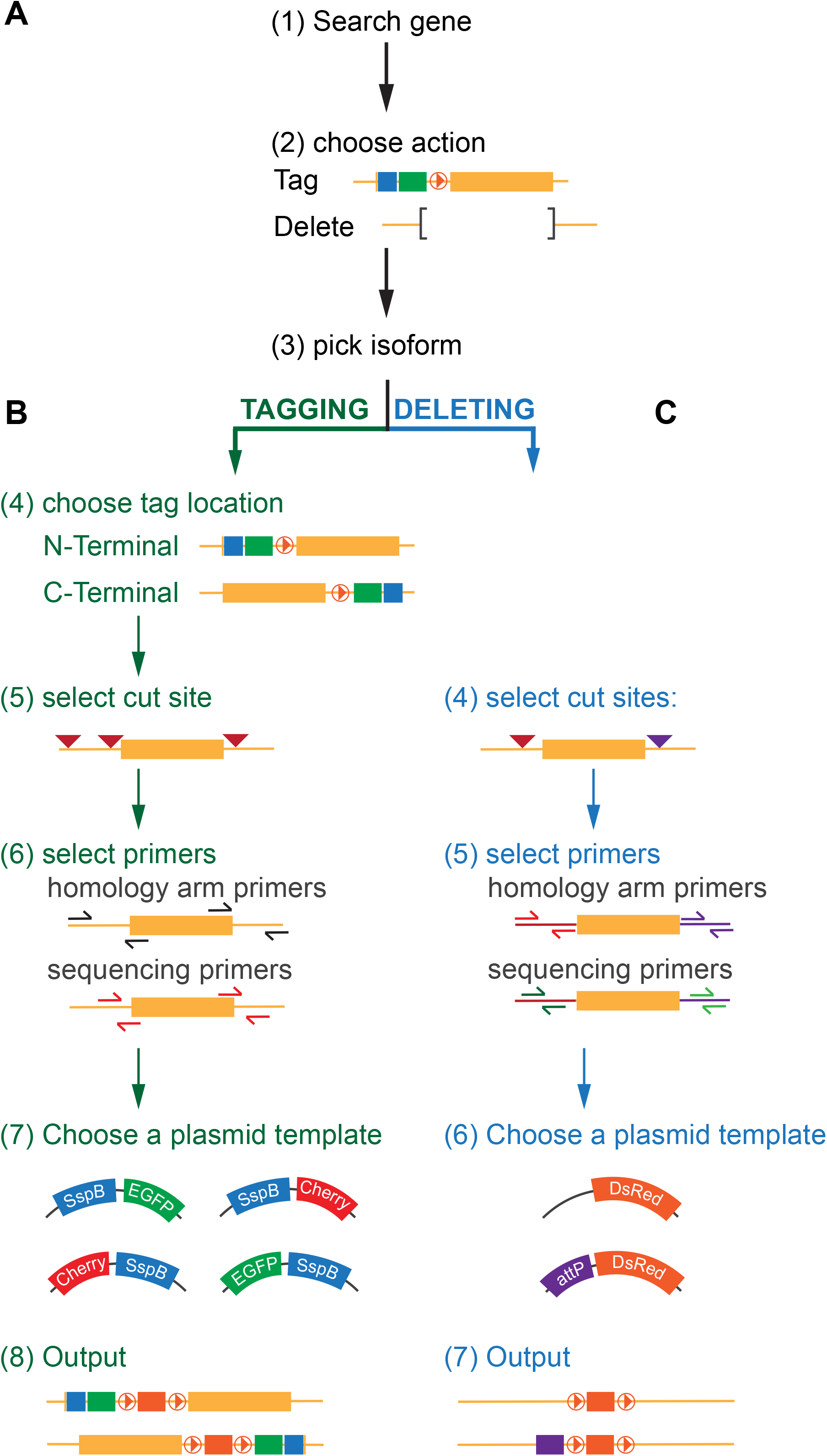
CrisprBuildr decision tree. CrisprBuildr, an open source genome engineering application allows users to design deletion and tagging experiments in silico. **(A)** After entering a *Drosophila* gene of interest, users can **(B)** tag or **(C)** genes of interest. The application prompts users to pick the desired gene isoform, suggests Cas9 cutting sites, proposes primers to amplify and sequence homology arms. Users can pick from a selection of tagging or deletion vectors, depending on the experiment. The application output are (1) a pU6-BbsI-chiRNA vector map containing the selected guide RNAs, (2) a unaltered map of the genomic locus and (3) a map containing the selected cassette in the desired location: SspB::EGFP/mCherry for N-terminal tagging, mCherry/EGFP::SspB for C-terminal tagging or attP-DsRed and DsRed, respectively, replacing the gene of interest.

To delete genes of interest the app is following a similar workflow, with the exception that two cutting sites are chosen. For instance, to delete genes or exons of genes, users can pick an N’ and C’ terminal cutting site. Subsequently, amplification and sequencing primers for both the N- and C-terminus are picked. Users can pick pHD-DsRed or pHD-DsRed-attP as integration vectors, replacing the targeted gene (Figure 3C).

For both operations the application provides a printable project information review and three downloadable cloning maps: (1) a pU6-BbsI-chiRNA vector map containing the selected guide RNAs to guide the Cas9 endonuclease into their target sites for proper double strand breakage; (2) A gene locus map without the chosen tag, containing the chosen gene isoforms and primers to amplify the homology arms as well as sequencing primers; (3) a gene locus map with the SspB::FP-DsRed, FP::SspB-DsRed, DsRed or DsRed-attP inserted at the desired site. This map also displays the locus with the amplified homology arms. Ideally, this map can be used as a reference map for the actual cloning steps, but users should keep in mind to mutate the PAM site and verify the reading-frame after LoxP floxing. These maps can be downloaded and viewed in Ape (https://jorgensen.biology.utah.edu/wayned/ape/). A short CrisprBuilder user manual can be found in the supplemental materials and is linked to the application.

## Discussion

CRISPR-based genome engineering opened the door for targeted genome alterations with unprecedented precision. However, designing the necessary vectors and constructs can be time-intensive and require the necessary expertise. Here, we report the generation of new vectors to tag genes at the endogenous locus with CRISPR in *Drosophila melanogaster*. These vectors allow the insertion of fluorescent proteins – EGFP or mCherry – combined with the short iLID-binding peptide SsrA at the N- or C-terminus of fly genes of interest. We also built a web-based application, called CrisprBuildr (http://142.93.118.6/) that aids users in the design of deletion or tagging experiments. The app links to existing databases that identify Cas9 cut sites, aid in primer design and retrieve gene locus information from existing databases. CrisprBuildr guides users through the process in a stepwise fashion and generates an unaltered map of the locus of interest as well as a map of the locus after the deletion or insertion. This map can provide a blueprint to aid users in the cloning process and can be used for effective teaching in the design and cloning of constructs for CRISPR-Cas9 experiments.

CrisprBuildr is ready to use but offers opportunities for future expansion. For instance, for gene-tagging experiments, we routinely mutate the PAM site in the donor construct to avoid Cas9 cutting in the repair template. In its current form, the application does not incorporate an automated way to mutate the PAM site, and users are advised to mutate it manually in the final cloning map and to design the cloning strategy accordingly. Similarly, only a small collection of deletion vectors (limited to pHD-DsRed-attB) or tagging vectors is included in the application but users could expand that choice by incorporating their own homebuilt genome engineering vectors. For instance, genes of interests could be tagged with HA, V5, FLAG or other small peptide tags. Alternatively, inserting transcriptional activators such as Gal4 (BRAND and Perrimon 1993), LexAop (Lai and Lee 2006) or QF (Potter et al. 2010) could generate new driver lines for misexpression systems.

Currently, CrisprBuildr is only retrieving genomic information for *Drosophila melanogaster* but since the application is completely open-source, we envision that future versions could incorporate genomes from other *Drosophila* species or other established or emerging model organisms.

## Methods

### Plasmid cloning

To generate vectors for N-terminal tagging, EGFP or mCherry (FP, hereafter) was first cloned into pHD-SspB-R73Q-DsRed utilizing restriction sites NdeI and Pmel to generate pHD-SspB-R73Q::FP-DsRed. An Mlul restriction site was inserted between the SspB and the FP to allow for easy swapping of the fluorophore for future applications. N-terminal homology arms corresponding to a gene of interest can be cloned into this backbone using restriction sites EcoR1 and NotI or NdeI. C-terminal homology arms can be inserted into AscI and Xhol for the C-terminal homology arm.

To generate C-terminally tagged vectors EGFP or mCherry (FP, hereafter) was inserted into pHD-SspB-R73Q-DsRed utilizing restriction sites Not1 and Mlul to generate pHD-FP::SspB-R73Q-DsRed, conserving both restriction sites for future fluorophore swapping. N-terminal homology arms corresponding to a gene of interest can be cloned into this backbone using restriction sites Srf1, EcoR1 and NotI. C-terminal homology arms can be inserted between and BgIII and XhoI All cloning steps were performed using InFusion cloning from Takara.

Both N- and C-terminal tagging vectors contain an extended flexible linker between SspB-R73Q and the FP.

The following final vectors were made:

*For N-terminal tagging:*

pHD-SspB-ExLK-EGFP-DsRed

pHD-SspB-ExLK-mCherry-DsRed

*For C-terminal tagging:*

pHD-EGFP-ExLK-SspB-DsRed

pHD-mCherry-ExLK-SspB-DsRed

### Live cell imaging

72 to 96 hours after egg laying, larval brains were dissected using microdissection scissors (Fine Science Tools catalog number 15003-08) (Segura and Cabernard 2023) and forceps (Dumont #5, Electron Microscopy Sciences, item number 0103-5-PO) in Schneider’s medium supplemented with 10% bovine growth serum (BGS) (HyClone, item number SH30541.03) and transferred to chambered slides (Ibidi, catalog number 80826) for imaging. Live samples were imaged with an Intelligent Imaging Innovations (3i) spinning disc confocal system, consisting of a Yokogawa CSU-W1 spinning disc unit and two Prime 95B Scientific CMOS cameras. A 60x/1.4NA oil immersion objective mounted on a both microscopes was used for imaging. Live imaging voxels are 0.22 × 0.22 × 1 μm (60x/1.4NA spinning disc).

### CrisprBuildr

CrisprBuildr connects to api.flybase.org and cross references the users search term with available data from flybase. If a relevant match is found, it will ping the flybase api in order to pull the raw genetic data. Targets around either the start or stop codon are retrieved from targetfinder.flycrispr.neuro.brown.edu/. Target efficiency is checked with www.flyrnai.org/evaluateCrispr/. The application will then look for primers within 400-600 bases and 1000-1200 bases both upstream and downstream from our target area. It will submit this information via simulated browser to bioinfo.ut.ee/primer3-0.4.0/ and will return the selection to the user. Once the user finished selecting their primers, the application will pull the relevant Oligo information from targetfinder.flycrispr.neuro.brown.edu/.

The source code can be found here: https://github.com/SelinaLiu8/fly-server

## Acknowledgements

We thank members of the Cabernard laboratory for testing CrisprBuilder as well as helpful discussions and comments. This work was supported by the National Institutes of Health (R35GM148160 to CC).

